# Pigmentation loci as markers for genome editing in the Chagas disease vector Rhodnius prolixus

**DOI:** 10.1101/2020.04.29.067934

**Authors:** M. Berni, D. Bressan, Y. Simão, A. Julio, P. L. Oliveira, A. Pane, H. Araujo

## Abstract

The kissing bug *Rhodnius prolixus* is a major vector for Chagas disease in the Americas, and also considered as the primary model for functional studies. Prospective transgenic approaches and genome editing strategies hold great promise for controlling insect populations as well as disease propagation. In this context, identifying visible genetic markers for transgenic methodologies is of paramount importance to advance the field. Here we have identified and analyzed the function of putative cuticle and eye color genes by investigating the effect of gene knockdown on fertility, viability, and the generation of visible phenotypes. Synthesis of the dark, yellow and tan pigments present in the cuticle of most insects depends on the function of key genes encoding enzymes in the tyrosine pathway. Knockdown of the *R. prolixus yellow* and *aaNAT/pro* orthologs produces striking alterations in cuticle color. Surprisingly, knockdown of *ebony* does not generate visible phenotypes. Since loss of *ebony* function results in a dark cuticle in several insect orders, we conclude that *R. prolixus* evolved alternative strategies for cuticle coloration, possibly including the loss of a pigmentation function for an entire branch of the tyrosine pathway. Knockdown of the *scarlet* and *brown* genes - encoding ABC transporters - alters cuticle and eye pigmentation, implying that the transport of pigment into proper organelles is an important process both for cuticle and eye coloration in this species. Therefore, this analysis identifies for the first time potential visible markers for transgenesis in a hemipteran vector for a debilitating human disease.

**Author Summary:** The hemipteran *Rhodnius prolixus* - also known as a kissing bug - is a main vector transmitting the parasite *Trypanosoma cruzi*, the causative agent of Chagas disease, a debilitating infection estimated to affect more than 6 million people in Central and South America. In order to limit disease spread, an important measure is insect vector control. However, kissing bugs - like other insects - develop resistance to insecticides. Alternative strategies based on transgenesis and the recently developed CRISPR-based genome edition hold great promise to control vector population or generate parasite-resistant insects. For these approaches to be feasible in *R. prolixus*, it is critical to identify visible phenotypic markers. Here we identify and describe several genes controlling cuticle and eye pigmentation that are well-suited putative landing sites for transformation strategies. Among these, loss-of-function mutations in the ABC transporter encoding *scarlet* and the tyrosine pathway enzyme encoding *aaNAT/pro* generate striking and easily visible phenotypes. Importantly, the knockdown of these genes does not affect insect viability and fertility under laboratory conditions. Our results suggest that *R. prolixus* has developed alternative strategies for cuticle coloration involving the loss of an entire branch of tanning loci, while the other branch producing cuticle patterns by generating non-pigmented areas has gained critical importance.

## Introduction

Pigmentation is largely recognized as an evolutionarily selected trait in insects and other metazoans, with important functions in mate choice, camouflage, thermoregulation and resistance to desiccation and infection, among others [1]. Mutations in genes implicated in pigment synthesis pathways were first identified in *Drosophila melanogaster*, such as the eye color mutant *white* (*w*, [2,3]) and cuticle color mutants like *yellow* (*y*, [4,5]). Orthologs encoding pigment pathway enzymes have also been identified in several insect orders aside from Dipterans. Among Hemiptera (true bugs), cuticle coloration has been functionally analyzed in three plant eating species: *Oncopeltus fasciatus* [6,7], *Acyrthosiphon pisum* [8] and *Nilaparvata lugens* [9]. However, the Hemiptera order also harbors a great number of blood feeding insects. Kissing bugs, including species belonging to the Triatoma, Panstrongylus and Rhodnius genera, display a range of cuticle and eye color patterns, yet the evolutionary conservation of the pigment pathway in these species has not been investigated.

*Yellow (y*) is part of a highly conserved modular gene network that controls melanin and sclerotin production for cuticle coloration and sclerotization (Fig 1A). This biosynthetic pathway starts with hydroxylation of phenylalanine to tyrosine, followed by hydroxylation to dihydroxyphenylalanine (DOPA) [10], by the enzymes phenylalanine hydroxylase (PAH) and tyrosine hydroxylase (TH), respectively. DOPA is the substrate for the DOPA black melanin branch, and is also converted to Dopamine by DOPA decarboxilase (DDC), the substrate for the Dopamine black/brown melanin branch. *y*, that encodes dopachrome conversion enzyme (DCE), participates in both pathways to oxidate the DOPA and Dopamine precursors, generating the dark melanin pigment within the cuticle [5,11–15]. The Dopamine precursor is also used by N-beta-analyldopamine hydroxilase (NBAD) to generate yellow/tan colored sclerotin pigment. The last branch requires dopamine N-acetyl transferases (NAT) to convert Dopamine to N-acetyldopamine (NADA), which is the precursor of unpigmented NADA sclerotin (Shamim et al, 2014). Phenoloxidases (POs) participate in all branches of the pigmentation pathway. The *tan, ebony* and *black* genes reported in *Drosophila* [16,17], the silkworm *Bombyx mori* [13], the butterfly *Vanessa cardui* [15] and the beetle *Tribolium castaneum* [18], encode NBAD pathway enzymes. *aaNAT*, with reported roles in pigment pattern of *O. fasciatus* [7] and *Bombyx mori* [19], encodes the central enzyme that generates unpigmented NADA sclerotin (arylalkylamine-N-acetyltransferase) (Fig 1A).

**Fig 1.**
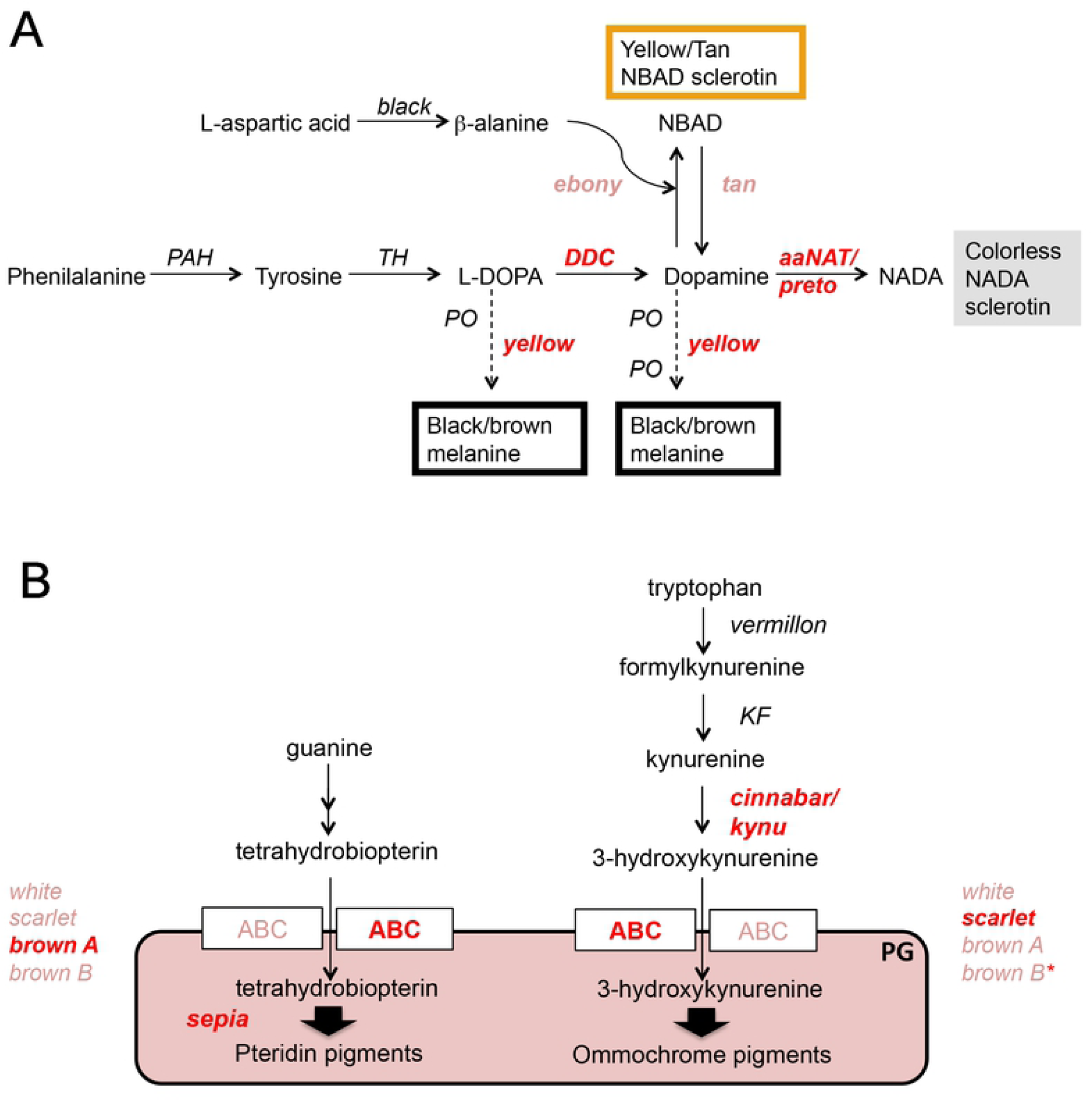
Schematics of pigmentation pathways and *Rhodnius prolixus* loci associated to cuticle and eye color. We have investigated cuticle pigmentation and eye color phenotypes resulting from knockdown for loci associated to: (A) the tyrosine pathway, involved in cuticle pigmentation in insects, and (B) pathways associated to the generation of ommochrome and pteridin pigments in the eye. In italic: red represents loci that resulted in a change in color compared to wild-type; pink indicates loci that resulted in no visible effect upon knockdown; black refers to loci that were investigated by others. PAH, phenylalanine hydroxylase; TH, tyrosine hydroxylase; DDC, DOPA decarboxylase; PO, phenoloxidase; KF, kynurenine formamidase. PG, pigment granule

In insects, eye color requires the synthesis of ommochrome and pteridine pigments from tryptophan and guanine precursors, respectively, and their uptake into pigment granules by ABC transporters [10,20] (Fig 1B). *vermillon* (*v*), that encodes tryptophan oxidase, and *cinnabar* (*cn*, or *kynu*), that encodes kynurenin hydroxylase, are associated with the production of ommochromes [21–24]. Loss-of-function mutants for *sepia* (*se*), that encodes a glutathione-S-transferase, are characterized by loss of the bright red eye color in *D. melanogaster* due to a decrease in red pteridins [25]. In *D. melanogaster* [26] and *Tribolium castaneum* [27] *w, scarlet* (*st*) and *brown (bw*) loci encode ABC transporters that for heteromeric channels: proteins encoded by *bw* and *w* transport the red pteridines into cells of developing eyes, while *w* plus *st* transport brown ommochrome pigments. Eye color mutations affecting components of either pathway have been described in several insect orders [9,27–34].

The cuticle pigment pathway is also important for melanization associated with the immune response [35,36] and for detoxification of tyrosine ingested from diet [37], which is of particular importance for insects feeding on blood. Due to the excessive amount of protein ingested during the blood meal, mounting to several times their body weight, insects depend on the detoxification of dietary tyrosine for optimal fitness. Therefore, it has been suggested that blood-feeding insects evolved strategies to reduce the redox stress employing the tyrosine pathway [38]. Thus, investigating how the melanin pathway was coopted to serve both for body color and a detox functions in blood feeding species may be particularly relevant for the biology of blood-sucking insects. A similar problem arises from the high levels of tryptophan present in the blood meal, also an essential precursor for ommochrome synthesis. Accordingly, the midgut of *R. prolixus* expresses high levels of both tryptophan and tyrosine degradation pathway enzymes after a blood meal [39].

The identification of loci that control pigmentation has lead to great advances in the development of transgenic strategies in insects. For instance, the identification and cloning of *D. melanogaster w*, and of *rosy*, that codes for the xanthine dehydrogenase required for pteridin synthesis in the eye, and the establishment of *w-* and *rosy-* mutant lines granted the development of transposable element-based transformation [40]. A decade later, identification of the *Ceratitis capitata w* gene enabled the development of germline transformation protocols in the medfly [41,42]. Visible markers were also decisive for modern CRISPR-based genome edition. In the first report that proved the principal of auto-replicative CRISPR-based gene drive elements in *D. melanogaster, y* was used as target for gene disruption [43]. Similarly, mosquito *kynurenine hydoroxilase* was used to reveal the feasibility of gene drive in *Anopheles stephensis* [44]. Notably, in the context of gene drive lineage production, pigmentation genes may both serve as visible markers carried on transgenic constructs as well as markers for gene disruption, when used as target sites for the incorporation of gene drive constructs.

As kissing bugs play a major role in transmission of parasitic Chagas disease, identifying cuticle and eye color loci will aid in defining visible markers for future transgenic and gene edition protocols. We hereby describe the identification and functional analyses of pigmentation pathway genes in *R. prolixus*. We show that knockdown of the *R. prolixus* orthologs of the *st* and *aaNAT* genes produce striking cuticle and eye color phenotypes. Remarkably, downregulation of these genes by RNA interference (RNAi) does not affect animal viability and female fertility in laboratory conditions. We demonstrate that several genes belonging to the cuticle color module display a conserved function in this insect, while one specific branch of the pathway, namely the NBAD tanning branch, appears to have lost most of its role in cuticle coloration. Our data shed light on the evolution of the pigmentation pathway in insects and provide easily scorable phenotypic markers, which will facilitate transgenesis, genome edition and, ultimately, the development of *R. prolixus* population control strategies.

## Results

### Identification of pigmentation genes in *R. prolixus*

We have identified several putative cuticle and eye pigmentation genes in the *R. prolixus* genome, based on protein sequence similarity to *D. melanogaster, O. fasciatus* and *Anopheles sp*. One clear ortholog was identified for each of the most evolutionarily conserved genes, particularly those coding for enzymes of the tyrosine, ommochrome and pteridine synthesis pathways (Table 1; S1 Table). The expression for many of these genes was confirmed by performing searches in transcriptomic datasets (data not shown). Differently, for the identification of *y* as well as of genes encoding putative ABC transporters, phylogenetic analyses were performed to identify the most likely orthologs among several sequences (S1 and S2 Fig). In agreement with the diversity of *y* paralogs reported in different insect species, we identified four *y* loci, three clear orthologs of *D. melanogaster y, y B* and *y C*. The forth *y*, which we termed *yellow-like* (*y-like)*, probably diverged from the main *y* branch. For eye pigment ABC transporters, we found that *R. prolixus* presents one *w*, one *st*, and two *bw* orthologs, which we refer to as *bw A* and *bw B*. This duplication event might be unique to *R. prolixus*, since only one *w*, one *st* and one *br* gene have been described in *D. melanogaster, T. castaneum [27]*, and *B. mori* [34]. Notably, the analysis presented below shows that most of the genes herein identified are functional, confirming our *in silico* identification.

**Table 1.** Pigmentation-associated genes identified and functionally analyzed in. this study. Genes were identified by sequence similarity to *D. melanogaster* and *Anopheles sp*. genes and validated by phylogenetic analysis. Gene names were defined following terminology used for other insects, mostly based on the loss-of-function phenotype.

| locus | symbol | protein/enzyme encoded | process involved | accession # |
| --- | --- | --- | --- | --- |
| *DDC* | *DDC* | Dopa decarboxylase | tyrosine pathway | RPRC005884-RA |
| *aaNAT/preto* | *pro* | Arylalkylamine-N-acetyltransferase | tyrosine pathway/NADA branch | RPRC015310-RA |
| *ebony* | *e* | N-beta-analyldopamine syntase | tyrosine pathway/NBAD branch | RPRC007578-RA |
| *tan* | *t* | N-beta-analyldopamine hydroxylase | tyrosine pathway/NBAD branch | RPRC007817-RA |
| *black* | *bl* | aminoacid decarboxiyase | tyrosine pathway/NBAD branch | RPRC010142-RA |
| *yellow* | *y* | Dopochrome conversion enzyme (DCE) | tyrosine pathway/ melanin synthesis | RPRC005424-RA |
| *yellow C* | *y C* | " | tyrosine pathway/ melanin synthesis | RPRC008209-RA |
| *yellow-like* | *y-like* | " | tyrosine pathway/ melanin synthesis | RPRC014337-RA |
| *sepia* | *se* | Gluthatione-S-transferase | pteridine synthesis | RPRC007741-RA |
| *white* | *w* | ABC transporter | eye pigment transporter | RPRC012709-RA |
| *scarlet* | *st* | " | eye pigment transporter | RPRC010854-RA |
| *brown A* | *bw A* | " | eye pigment transporter | RPRC009214-RA |
| *brown B* | *bw B* | " | eye pigment transporter | RPRC002598-RA |
| *cinnabar* | *cn* | Kynurenine hydroxylase | ommochrome synthesis | RPRC001714-RA |

### Visible phenotypes associated with loss of function for genes in the melanin synthesis pathway

To identify suitable loci as visible markers for transgenesis, we initially investigated the effect of tyrosine pathway genes on viability of the progeny and putative changes in cuticle pigmentation (Fig. 2; S2 Table). To this aim we performed parental RNAi (pRNAi) by injecting double-stranded RNA (dsRNA) molecules specific to each cognate gene into the hemocoel of adult females. A decrease in adult or embryo viability upon gene knockdown (KD) would exclude the gene as a good landing site for transgenesis. This was particularly true for *DDC*, even though the few surviving *DDC* KD embryos displayed total loss of pigmentation, as previously reported (S2 Table; [45]). On the other hand, *y* KD (RPRC005424) displayed a change in cuticle coloration in the thorax, head and legs, with no significant decrease in viability (Fig 2C,D). Additional tyrosine pathway loci presented no effect on pigmentation by pRNAi. We also investigated whether the blood diet had any effect on animals resulting from pRNAi. We observed no effect beyond those already present at nymph eclosion.

**Table 2. General phenotypic effects of tyrosine, ommochrome and pteridine pathway gene knockdowns**

**Fig 2.**
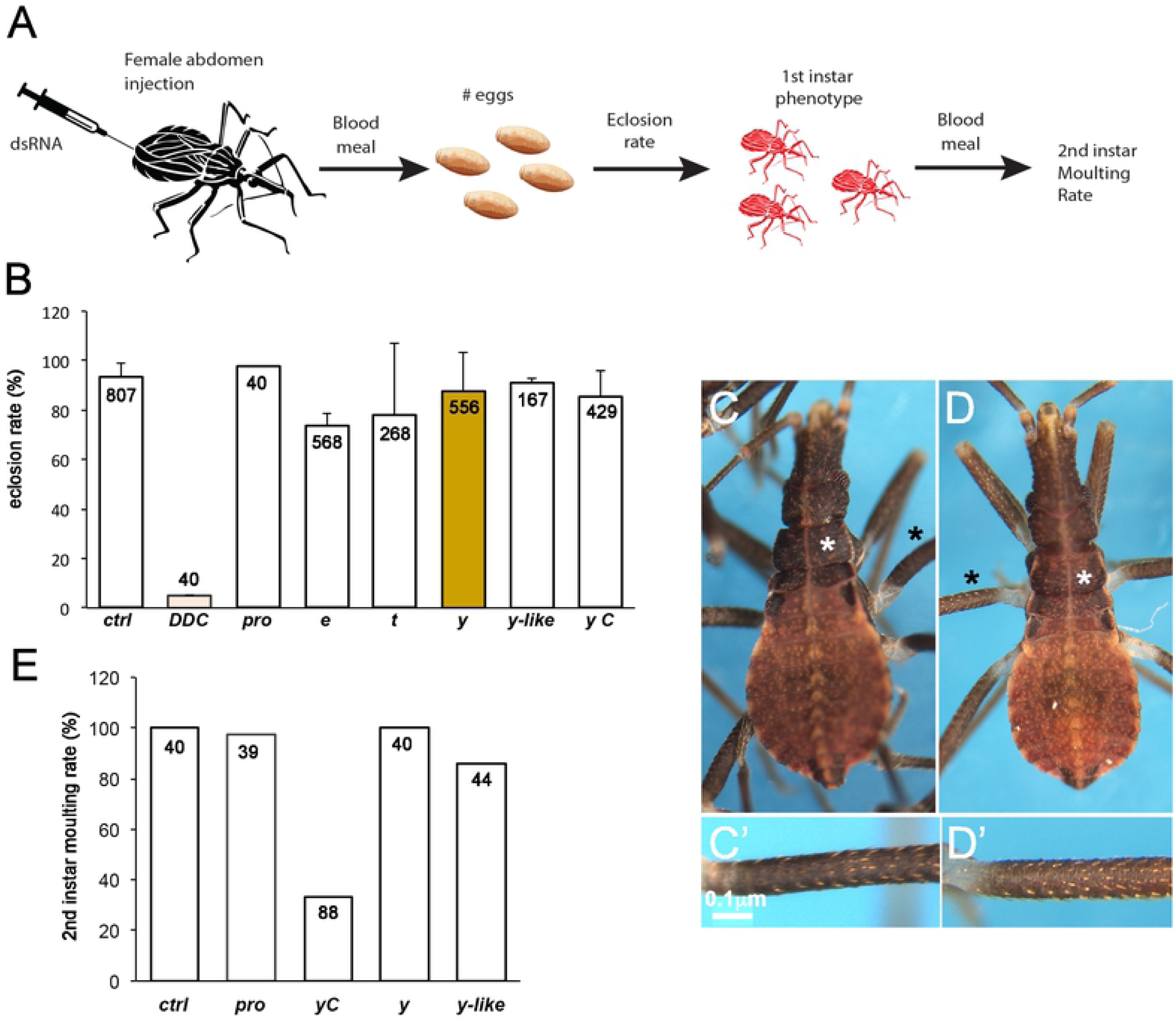
Parental knockdown for a limited set of putative *R. prolixus* tyrosine pathway loci affects embryo and first instar viability. We performed pRNAi for tyrosine pathway loci and analyzed their effect on injected animals and their progeny. A) Injection protocol for pRNAi. B) Viable progeny resulting from dsRNA injected females. C,D) First instar nymphs resulting from females injected with control (C) or *y* (D) dsRNA, showing that *y* KD animals have a slightly lighter cuticle, especially visible in the thorax and legs (asterisks). E) Molting rate to second instar of first instar nymph offspring from dsRNA injected females, after a blood meal. Only animals from *y C* dsRNA-injected females showed significant loss in viability.. Numbers displayed inside bars correspond to individual eggs (B) or first instar nymphs (E) analyzed.

Next, we assayed tyrosine pathway gene function by injecting dsRNA in fifth instar nymphs. After feeding, these nymphs molt in around 16 days as unpigmented adults, regaining full color in approximately 12 hours. In this condition we observed a significant change in cuticle color for *y* and *aaNAT* KD (Fig 3). This is especially evident in the thorax, where three different colored stripes are seen: dark stripes which we called black (bl), brown stripes we termed tan (tan), and clear stripes identified as white (wh). In *y* KD the bl stripes are lighter, consistent with a role for *y* in generating black/brown melanin (Fig 3C). Two other *yellow* loci, *y C* (RPRC008209) and *y-like* (RPRC014337) displayed only a weak effect on cuticle pigmentation. However, the *y C* plus *y-like* double KD resembles the *y* KD cuticles, suggesting redundancy among *yellow* paralogs (S3 Fig). On the other hand, *aaNAT* KD cuticles display a striking phenotype: they are homogeneously dark, loosing the tan and wh stripes of the thorax and any pattern characteristic of the insect throughout the entire body (Fig 3E). Since *aaNAT* is predicted to generate uncolored sclerotin, this observation indicates that *aaNAT* function is required to produce all the light color patterns of the insect body, including the tan and wh stripes of the thorax. Such effect is surprising given the weak effect of *aaNAT* loss-of-function shown in other insects, where only a few dark spots are gained [7,19]. Equally surprising is the fact that *ebony* (*e*) and *tan* (*t*) KD had no effect on pigmentation, given the extremely dark loss-of-function phenotype observed in several insect species (S2 Table; [7,13,15,46]). Unfortunately, we were unable to obtain a *black* cDNA for dsRNA production and functional analysis using ovaries or embryos. Low expression levels in the gut and early embryogenesis suggest, however, that *black* is functional in other tissues. Due to the strong phenotype displayed by *aaNAT* KD, we henceforth refer to the locus associated to this function as *preto* (*pro*), the Brazilian portuguese term for “black”.

**Fig 3.**
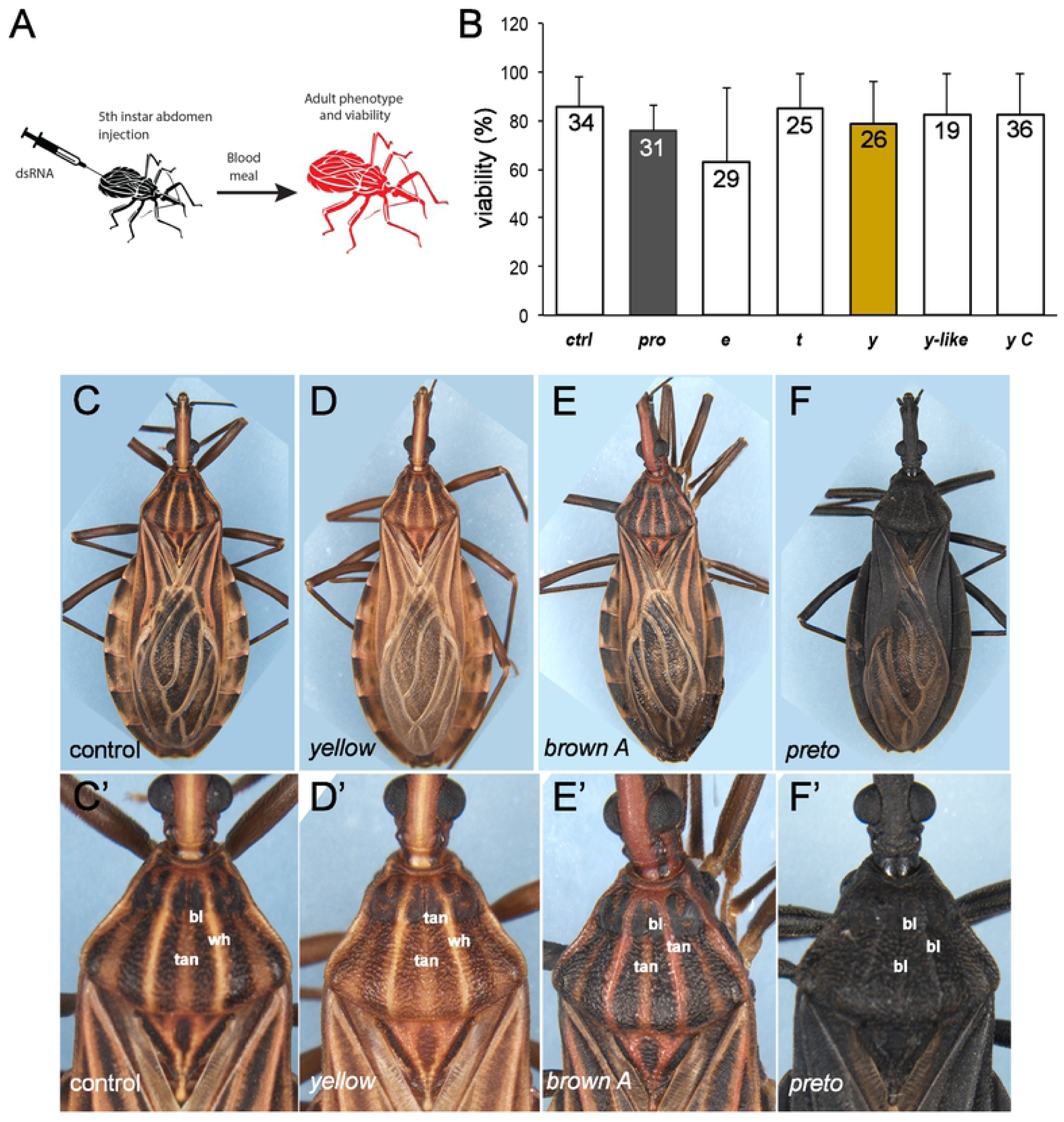
Cuticle pigmentation phenotypes resulting from knockdown of the tyrosine pathway genes. A) RNAi strategy: fifth instar nymphs were injected with dsRNA specific to tyrosine pathway genes and the phenotype was analyzed in the adult. B) Viability of animals injected with dsRNA, after a blood meal. No significant difference between experimental conditions versus control was observed Numbers in histograms correspond to the number of fifth instar nymphs injected. C-F) Control (C,C’) and cuticle phenotypes resulting from knockdown for *y* (D,D’) *bw A* (E,E’), and *aaNAT/pro* (F,F’). C’-F’) Higher magnification of C-F showing details of the three-color pattern of the *R. prolixus* first thoracic segment in control and knockdown animals.

To further investigate the effect of different melanin pathway branches on cuticle pigmentation, we performed double knockdowns for *e* plus *y* as well as for *pro* plus *y* (Fig 4). Double KD for *e* and *y* produced a phenotype comparable to *y* KD alone, both with respect to head, thorax and abdomen patterns as well as to wing pigmentation (Fig 4A-C, G-J). Together with the single KD assays reported above, this observation supports the conclusion that the NBAD branch does not control cuticle pigmentation in *R. prolixus*. On the other hand, *pro* plus *y* double KD generates animals that are unpatterned and light brown up to 24h after molting (Fig 4D-F, K-L), when control insects are already fully pigmented. Subsequently, these animals gain a dark cuticle that resembles the *pro* KD phenotype. This delayed pigmentation pattern observed in the double *y* + *pro* KD could be explained by a decrease in enzymatic activity in both the NADA (*pro*) and brown/black melanin (*y*) branch, with residual Yellow activity slowly building up the dark pigmentation.

**Fig 4.**
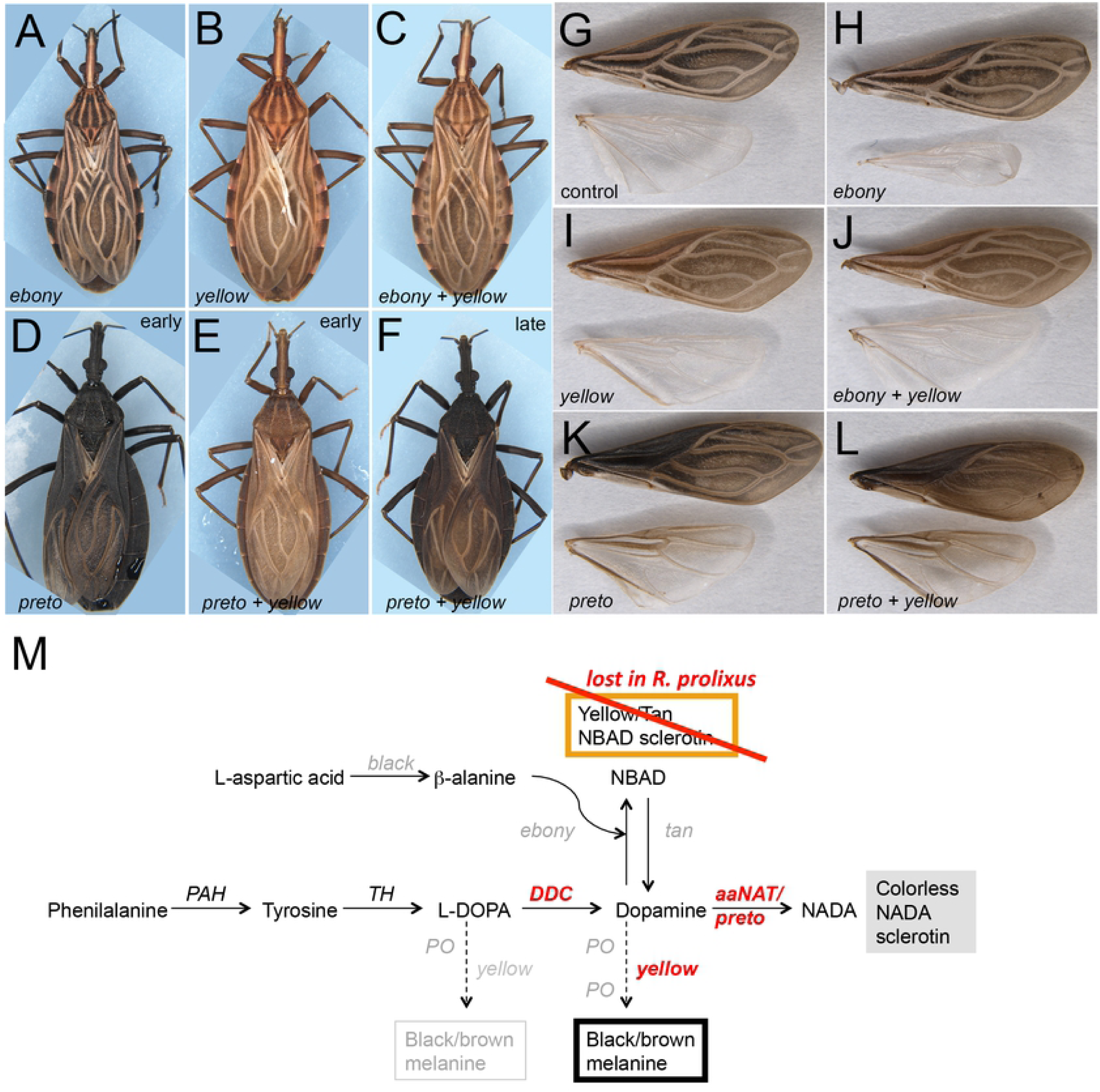
Double knockdown for distinct tyrosine pathway branches points to the absence of a cuticle pigmentation function for the NBAD branch. Effects on body (A-F) and wing (G-L) pigmentation, resulting from single or double KD for tyrosine pathway loci. A,H) *e* KD cuticles are identical to wild-type or control (G); B,I) *y* KD; C,J) *e + y* KD; D,K) *pro* KD; E) *y* + *pro* KD at 24 hours after moulting; F,L) *y* + *pro* KD at 48h hours after moulting. G) Wings from control KD.

### Loss of function for classical eye color genes generates visible phenotypes

*Anopheles stephensi kynu*, the ortholog of *D. melanogaster cn*, has been successfully used as a phenotypic marker in previous transgenic studies, since loss of *kynu* function generates white eyes [44,47]. In order to investigate the function of the *R. prolixus cn/kynu* ortholog, we injected dsRNA in fifth instar nymphs, and looked for a visible eye color phenotype in the adults that emerge after molting. As a result, *cn* KD adult *R. prolixus* have reddish eyes (Fig 5A,B,D). pRNAi results in progeny displaying a similar eye color phenotype. These first instar nymphs have either reddish eyes or display a red circle around the black colored eye (Fig 5F,G,H). Interestingly, this pattern is observed in early molting wild-type animals where the wild-type black color slowly builds from a red eye, but, unlike *cn* KD, the red circle disappears with age (S4 Fig). Importantly, we observed no effect of *cn* KD on viability of the egg/first instar nymph before a blood meal. However, feeding *cn* KD first instars with a blood meal resulted in death of all animals before they could molt to second instars (Fig 5J). This suggests that *cn* function is essential for the molting process.

**Fig. 5.**
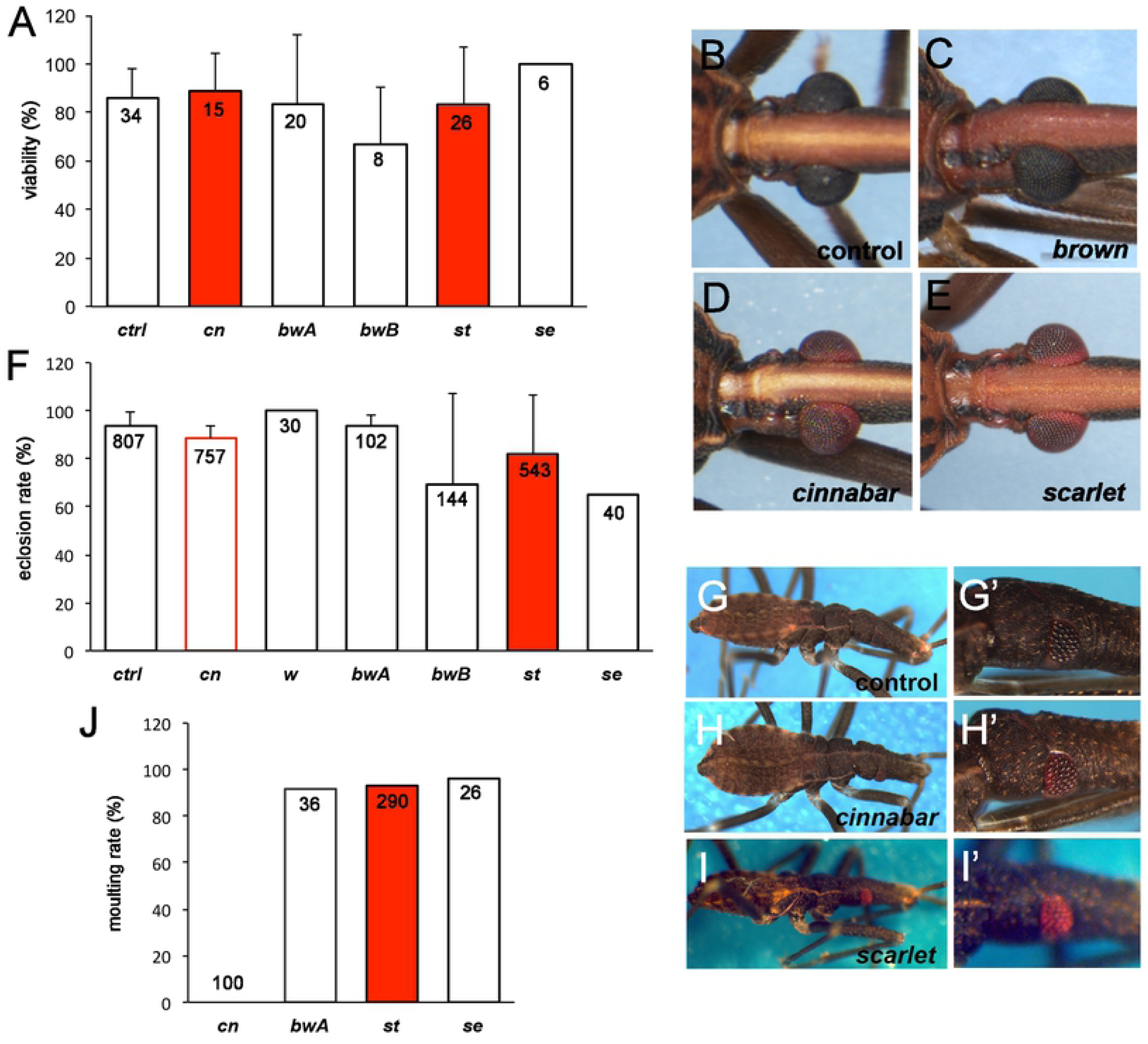
Phenotypes resulting from the knockdown of genes involved in the production and transport of eye pigments. A) Adult viability and B-E) Adult eye color phenotypes observed following fifth instar dsRNA injections for B) control, C) *bw A*, D) *cn* and E) *st*. Numbers in histograms correspond to the number of fifth instar nymphs injected. F-I) First instar phenotypes resulting from adult female dsRNA injections. F) Egg viability resulting from pRNAi. G-I) First instar eye color phenotypes observed following pRNAi for G,G’) control, H,H’) *cn* and I,I) *st*. J) Molting rate to second instar of first instar nymph offspring from dsRNA injected females after a blood meal. The number of individual first instar nymphs analyzed is displayed inside bars. Note that after blood ingestion none of the *cn* KD animals survived.

Next, we performed KD for putative ABC pigment transporters. Surprisingly, the KD of the *w* and *bw B* orthologs in *R. prolixus* did not produce obvious phenotypes (Supp Table II), although red pigment spots in adult connexives are lost in *bw B* KD (Fig S3). Conversely, KDs for *st* and *bw A* showed striking effects (Fig 5): *st* KD gives rise to red eyes, suggesting that *st* acts in the eye for the transport of dark ommochrome pigments (Fig 5E,G,I). *st* KD resulting from dsRNA injection in fifth instar nymphs also alters cuticle coloration in addition to generating a red eye phenotype. Cuticles are reddish in appearance, missing the wh stripes of the head and body (Fig 5B,E). On the other hand, *bw A* has no effect on the eye, but shows a clear effect on the cuticle (Fig 5B,C). In the animal’s thorax, originally light regions of the cuticle are now tainted red as shown for *st* KD (Fig 5C and 3E). Interestingly, KD of *se*, whose protein product has been reported to function in the synthesis of pteridines in several insect species, generates a thoracic pigmentation phenotype that is identical to *bw A* KD (S3 Fig). This suggests that *bw A* and *se* KD result in loss of a pigment that acts to mask the red cuticle color. Alternatively, red pteridines are transported away from the cuticle or modified to another color in order to maintain light stripes and patterns.

## Discussion

### An alternative strategy for cuticle pigmentation in *R. prolixus*

We have analyzed several *R. prolixus* loci coding for enzymes of the tyrosine pathway, which have been classically associated to the production of cuticle color. Genetic loci that encode enzymes in the initial steps of the tyrosine pathway frequently produce changes in cuticle coloration as well as cuticle integrity/strength, consistent with branching of the pathway for the production of melanin and sclerotin. Accordingly, knockdowns and loss-of-function mutants for the genes encoding the initial enzymes of the pathway PAH, TH and DDC lead to loss of cuticle coloration in *D. melanogaster, O. fasciatus* and *V. cardui*, to soft cuticles in *T. castaneum* and *Anopheles sinenses* [14,48], and decrease in first instar hatching in *R. prolixus* and *B. mori* [38,49]. The reduction in hatching rates may result from softness of the eggshell, as eggshell structure depends on adequate sclerotization, making it difficult for the emerging nymphs to punch through the eggshell. Furthermore, pigmentation may be associated to desiccation resistance as shown for *Aedes, Anopheles* and *Culex* [50].

The function of phenoloxidases (PO), which participate in all melanin and sclerotin branches, was previously analyzed in *R. prolixus* by gene knockdown and did not produce apparent phenotypes [37]. However, due to the large number of PO coding genes (at least four) in the *R. prolixus* genome, this may result from functional redundancy. Subsequent steps of the tyrosine pathway are restricted either to the melanin or sclerotin production pathways, and are thus more likely to contribute useful phenotypic markers. Among the putative loci tested, *y* produced the expected yellow loss-of-function cuticle phenotype, where dark black stripes of the adult thorax are lightened and wings lose their dark coloration. This is consistent with a role in generating brown/black melanin. Furthermore, the strong effect of the double *y C plus y-like* KD, as compared to single KDs, suggests functional redundancy of *yellow* loci. Phylogenetic analysis of *R. prolixus yellow* genes conforms to this view (S2 Fig).

Using Dopamine as substrate, the NBAD sclerotin branch has been associated to the generation of tanned/yellow cuticle. In *Drosophila, Oncopeltus, Tribolium, Bombyx* and *V. cardui, e* (*ebony*) and *bl* (*black*) KDs produce animals with dark cuticle due to loss of dopamine conversion to the tan/yellow pigment, leaving Dopamine available for the conversion to black pigment [7,13,15,46,51]. Conversely, loss of *t* (*tan*) function results in light pigmented cuticles in *Drosophila, O. fasciatus, N. lugens* and *C. capitata* [7,46,52]. The functions of the *e* and *t* genes in blood feeding insects has not been investigated yet, although a study in *A. gambiae* showed that a *bl* allele generates animals with dark cuticle and reduced fertility and vigor [53]. Here we show that *R. prolixus e* and *t* KD do not produce a visible phenotype, although a small reduction in insect viability was observed in both assays and mRNA levels clearly reduce (S5 Fig). NBAD synthase and NBAD hydrolase, encoded by *e* and *t*, respectively, have been implicated in several different processes. They exert a role in brain neurotransmitter metabolism [52], and *e* is expressed in the foregut and tracheal epidermis in *D. melanogaster* [51]. Their effect on tyrosine detoxification, which could also explain the loss of viability, has not been explored.

Surprisingly, inhibiting the NADA branch by *aaNAT*/*pro* KD results in a much stronger phenotype than shown for other insects [7,19]. The dramatic effect of *pro* KD on cuticle pigmentation in *R. prolixus* suggests that AaNAT activity is required to produce low-pigmented areas and thus generate specific cuticle patterns by “erasing” color. Taking into account the lack of any visible effect of *e* and *t* KDs, this suggests that *R. prolixus* evolved the preferential use of non-pigmented NADA sclerotin and dark melanin as pigments for cuticle coloration pattern, with little or no contribution from the NBAD synthesis enzymes (see Fig. 1). Recently, it was shown that *aaNAT* RNAi in the black colored *Platymeris biguttatus* bug obliterates white spots as well as yellow and red colors. Together with *aaNAT/pro* KD phenotype in *R. prolixus*, these findings may suggest that hemimetabolous blood feeding insects rely greatly on the NADA branch for cuticle color patterning.

Notably, kissing bugs display dark and frequently monotonous pigmentation patterns, as compared to their plant feeding relatives [54]. Among blood feeding Triatomines, the sole color that diverges from the black-tan-clear pallet is red. Interestingly, the red color originates in *R. prolixus* head and thorax as a result of *st, bw A* and *se* KDs (Fig 3E, Fig 5C and E; S3D Fig), resembling some *Triatoma* and *Panstrongylus* species cuticle pattern. Differently, *bw B* KD results in loss of red pigments that are brought through small veins that connect to the abdominal connexivum (S3F Fig). This indicates that red pigments, unrelated to the melanin pathway, are transported by ABC proteins to define the color of the kissing bug cuticle. In butterflies, the classical eye color-associated genes *v, cn*, and *w* are required for pigment patterning during wing development [55]. Importantly, ommochromes, pteridines and ABC transporters regulate cuticle pigmentation in many insect orders [10]. It will be interesting to investigate whether kissing bugs in general use a limited set of tyrosine pathway genes in addition to pteridine and ommochrome pigments for cuticle pigmentation as our results suggest for *R. prolixus*.

### Pigment transporters in *R. prolixus*

Loss-of-function of *w* genes generates eye pigmentation phenotypes in most insects studied to date. However, we observed no visible phenotype upon *R. prolixus w* KD. KD of *st*, on the other hand, generated red eyes and red tainted cuticle in the head and thorax. The red eye phenotype is likely due to a reduction in the transport of black ommochrome pigments, since KD for *cn*, that encodes an enzyme at the basis of the ommochrome synthesis pathway, generates a similar phenotype. An open question thus concerns the identity of the gene coding the ABC transporter that combines with the *st* product to form heterodimeric channels in the eye pigment granules, since we observed no eye pigmentation phenotype in KDs for other ABC transporters herein analyzed. In the head and thorax, St and Bw A likely interact to transport pteridine pigments, since *st, bw A* and *se* KDs display a similar phenotype. Notably, *bw B* KD results in loss of spatially restricted abdominal red spots, a phenotype only mirrored in the *cn* KD. Thus *bw B* probably transports ommochrome pigments in the abdomen. These results also suggest that *bw A* and *bw B* may have spatially distinct expression patterns, and transport different pigments in the head and thorax versus abdomen. In addition, *bw B* probably forms heterodimers with a different ABC protein than *st*, since *st* KD does not change the abdominal pattern. The lack of an eye color phenotype upon *w, bw A* and *bw B* KD is surprising, given the classical phenotypes reported for *D. melanogaster*. However, knockdown of a *T. castaneum brown* ortholog indicated no function in eye either [27]. Future studies directed to investigate functional redundancies in the genes coding for ABC transporters in *R. prolixus* may shed light on the control of eye pigmentation in this species.

### Visible markers for construct integration in *R. prolixus*

Using knockdown assays we have identified several loci that change eye or cuticle color and thus may serve as transformation markers in transgenic constructs or markers for construct integration upon CRISPR-based genome edition and HDR. Our results show that loss-of-function for a few loci generates visible phenotypes while having little effect on viability/fertility. Considering that the knockdown approximates the effect of gene disruption, these loci will certainly be useful as target integration sites. The most promising in this sense are *aaNAT/pro* and *st*, which present easily scored phenotypes and no significant effect on viability. *se* and *bw A* also share similar characteristics, although the red coloration is harder to perceive at a glance.

In contrast to the loci above, *y C* KD led to a small drop in nymph viability after blood feeding. This effect may result from loss of a waterproofing function, as shown for the beetle *T. castaneum* [56]. Therefore, despite the clear cuticle phenotype, apparent redundancy among *R. prolixus y* loci and broad use as marker in several insect species, we consider its potential as target integration site smaller than *pro* and *st*. Hence, our analysis has pointed out specific loci as markers for use in transgenic studies and genome edition in the kissing bug model *Rhodnius prolixus*.

## Materials and Methods

### Insect rearing

*R. prolixus* rearing was performed at 28°C and 70-75% humidity. Animal care and experimental protocols were conducted following guidelines of the Committee for Evaluation of Animal Use for Research from the Federal University of Rio de Janeiro (CAUAP-UFRJ) and the NIH Guide for Care and Use of Laboratory Animals (ISBN0-309-05377-3). Technicians dedicated to the animal facility at the Institute of Medical Biochemistry (UFRJ) conducted all aspects related to rabbit husbandry under strict guidelines to ensure careful and consistent animal handling.

### Identification of cuticle and eye color related genes in *R. prolixus* genome and phylogenetic construction

*D. melanogaster and Anopheles stephensi* protein sequences were used as query to BLAST into the *Rhodnius prolixus* genome (https://www.vectorbase.org/). After manual curation, protein sequences were aligned using the CLUSTALW algorithm available at the MEGA6 package [57]. Accession numbers for the genes analyzed are provided in Table 1. For phylogenetic analysis of ABC transporter and yellow genes, the evolutionary histories were inferred applying a Maximum Likelihood method [58] as described in Brito et al (2018)[59]. Briefly, the amino acid sequences were aligned by the Multiple Sequence Alignment with Log Expectation (MUSCLE, version 3.8.31) method [60], employing standard parameters. The evolutionary history was inferred by Molecular Evolutionary Genetics Analysis version 6.0 (MEGA6) [61], and visualized using interactive Tree of Life (iTOL, v2) [62]. The tree was validated by 500 bootstraps replications. All values higher of bootstraps were indicated in nodes. The amino acid sequences of proteins used in this study were obtained from VectorBase (https://www.vectorbase.org/), FlyBase (http://flybase.org/) and NCBI (https://www.ncbi.nlm.nih.gov) the identification of each protein is indicated in the tree.

### RNA interference assays (RNAi)

Double stranded RNA was synthesized from PCR products containing T7 promoter sequences at both ends as previously described [63]. Two successive PCRs were performed, the first to amplify the open reading frame of the gene of interest and the second added T7 promoter sequences at both ends. Primer pairs used in the first PCR are listed in Supp Table I. *In vitro* transcription was performed with Megascript kit (Ambion) as per manufacturer instructions. Two microliters of each dsRNA (1μg/μl) was injected in the abdomen of adult females three to five days prior blood feeding. Eggs were collected, counted, and the hatch rate defined after 20 days at 28°C. For fifth instar RNAi, 2 microliters dsRNA (1μg/μl) were injected into female or male abdomen. The insects were blood fed 5 days after the injection and let develop to the adult stage.

### Total RNA extraction and RT-PCR assays

For cDNA generation total RNA was extracted from eggs, first instar nymphs and carcasses using Trizol Reagent (Invitrogen) as per manufacturer instructions. Total RNA was treated with Turbo DNA Free (Ambion) to remove genomic DNA traces. The resulting DNA-free total RNA was subjected to *in vitro* Reverse Transcription (RT) with Superscript III (Invitrogen). 1µg of total RNA was used for each reaction and assays were conducted in biological triplicates. The oligonucleotides used in RT-PCR assays are listed in Suppl. Table I.

### Image processing

Microscopic images were obtained using a Leica Stereomicroscope, always on live animals. To minimize possible variation of the captured images, the background microscope and camera settings, as well image processing were standardized. To avoid cuticle pigmentation age related changes, all images were acquired during equivalent periods after molting. Adults were imaged 5 days after molt and first instars were imaged 2 days after eclosion.

## Acknowledgments

We would like to thank Dr. Annabel Guichard and members of the Araujo lab for helpful comments on the manuscript and Dr, Katia Gondim for great suggestions on Rhodnius husbandry. We also thank the insect facility at the Institute of Medical Biochemistry for maintaining the Rhodnius colony and providing healthy animals.

## Supporting Information Legends

**S1 Fig. Phylogeneticanalysisof *R. prolixusyellow*genefamily**. Genes were identified and names were given based on greatest similarity of R. *prolixus yellow* to previously characterized insect *yellow* genes.

**S2 Fig. Phylogeneticanalysisof *R. prolixus* ABCfamily transporters**. A limited set of *R. prolixus* ABC transporters is displayed, particularly those that displayed highest similarity to genes associated to pigmentation phenotypes in other insects.

**S3 Fig. Cuticle pigmentation resulting from knockdown for additional tyrosine pathway and pteridine synthesis loci**. Fifth instar nymphs were injected with 2μg/ml dsRNA and the phenotype was analyzed in adults. A) Recently emerged wild type adult before pigment deposition in the cuticle. Eyes are already dark. (B-J) 76h old adults resulting from dsRNA injections against B) *GFP*; C) *pugilist*; D) *sepia*; E) *cinnabar*; F) *brown B*; G) *tan*; H) *yellow C*; I) *yellow-like*; J) *yellow C-like* plus *yellow-like* (1μg/ml each). Upper right insets show eye color phenotype in *cinnabar* KD (E) compared to control (B). Bottom right insets show details of connexive pigmentation, with loss of red pigment spots in *cinnabar* (E) and *brown B* (F), compared to control (B) KD. Note the red cuticle color in *sepia* KD and the light color in *yellow C* plus *yellow-like* double KD.

**S4 Fig. Temporal progression of eye and cuticle color**. Color upon emergence of first instar nymphs (top) and emergence of adult male and female *R. prolixus* (bottom). The progressive appearance of a clear red ring around the eye and darkening with early aging implies that effects on eye color shoud be considered only after a 76 hour period, after which eye color is constant. For adults, the black eye color is already evident upon emergence, different from the cuticle that is initially uncolored and reveals the underlying pink tissues, and progressively darkens to full color at 24 hours after emergence.

**S5 Fig. Extent of gene knockdown for loci associated to cuticle and eye color phenotypes**. qRT-PCR analysis of recently emerged adult cuticles from fifth instar nymphs injected with dsRNA for (A) *yellow*, (B) *yellow-C*, (C) *ebony*, (D*) tan*, and (E) *scarlet*. Despite no visible phenotype, the levels of *ebony* and *tan* mRNAs show a significant decrease compared to wild type.

**S1 Table. Loci investigated in this study, with accession numbers and primers used for RNA interference**.

**S2 Table. Consolidated effects of gene knockdown for tyrosine (top) and tryptophan (bottom) pathway associated loci**. “Unpigmented connexives” refers to lack of red pigment in veins that transport pigments to connexives (S3 Figure).

